# Increased tolerance to commonly used antibiotics in a *Pseudomonas aeruginosa ex vivo* porcine keratitis model

**DOI:** 10.1101/2023.08.10.552790

**Authors:** K. Okurowska, P. N. Monk, E. Karunakaran

## Abstract

Antibiotics in development are usually tested on rapidly dividing cells in a culture medium and do not reflect the complexity of infections *in vivo*, while testing *in vivo* is limited, expensive and ethically concerning. This often results in the development and subsequent prescription of antibiotics only targeting infections in which pathogens are undergoing rapid cell division and in case of persistent infections like keratitis leads to poor clinical outcomes such as impaired vision or loss of an eye. In this study, we demonstrate antibiotic tolerance of *Pseudomonas aeruginosa* strains PA01 and PA14 using the *ex vivo* porcine keratitis model in which bacterial physiology more closely mimics infections *in vivo* than in a culture medium.

MBEC and MIC were used as a guideline to establish the concentration of applied antibiotics on tissue. Infected *ex vivo* porcine corneas were treated with therapeutically relevant concentrations of gentamicin, ciprofloxacin and chloramphenicol. Ciprofloxacin was the most potent across all tests demonstrating a positive correlation with MIC but not MBEC. Nonetheless, the results demonstrated that MIC and MBEC concentrations were not sufficient to clear infection even after 18 hours of continuous exposure to the tested antibiotics reflecting the need for novel antibiotics that can target the persistent subpopulation of these pathogens and the ability of the *ex vivo* keratitis model to be a relevant platform to identify novel antibiotics with suitable activities. There was a clear visual distinction between corneas infected with cytotoxic strain PA14 and invasive strain PA01. In this study, both strains PA14 and PA01 showed a high level of antibiotic tolerance, which suggests that in clinical settings the treatment approach could be similar regardless of the causative strain.

**Data summary:** The authors confirm all supporting data and protocols have been provided within the article or through supplementary data files.

## Introduction

Bacterial keratitis usually occurs because of infection following the trauma to the corneal epithelium caused by injury. Globally blindness caused by bacterial keratitis affects 1.5 to 2 million people each year (Whitcher et al., 2001, Humphries et al., 2019) although it is widely acknowledged that keratitis cases globally are underreported (Ung et al., 2019). Amongst the many pathogens that can cause bacterial keratitis, *Pseudomonas aeruginosa* is particularly difficult to treat and is a leading cause of sight loss in the developing world. Widespread use of antibiotics in livestock, availability of antimicrobial treatments without prescription and inappropriate prophylactic use contribute to higher antimicrobial resistance amongst these pathogens (Ting et al., 2021b, Hilliam et al., 2020, Willcox, 2011). Additionally, it is well known that *P. aeruginosa* forms biofilms. Extreme multi-drug resistance and poor clinical outcomes are hallmarks of biofilm infections (Maurice et al., 2018, Thi et al., 2020). Clinical isolates of *P. aeruginosa* resistant to the most used antibiotics are frequently found around the world (Lopez-Dupla et al., 2009, Garg et al., 1999, Willcox, 2011) and reinforce the global urgency to develop new antibiotics.

Administration of antibiotics in the early stages of infection is recognised clinically as essential for therapeutic success (O’Brien, 2003). Therefore, keratitis is considered as an ocular emergency and treated empirically with broad-spectrum antibiotics. Patients are usually prescribed fluoroquinolone monotherapy (e.g. ciprofloxacin) or a combination therapy with fortified antibiotics (Gokhale, 2008, O’Brien, 2003). In a few cases, prior to empirical antibiotic treatment, the corneal scrape is cultured to isolate the causative organism and antibiotic sensitivity testing is performed to select a more targeted (evidence-based) treatment. However, this is time-consuming, and growth and identification of microorganisms occurs in only 40-60% of cases. Evidence-based prescription of appropriate antibiotics is not routinely undertaken in clinics (Dalmon et al., 2012, Ibrahim et al., 2009, Norina et al., 2008, Varaprasathan et al., 2004).

Currently, treatments aim to achieve the minimum inhibitory concentration (MIC) of the drug at the site of infection (Gokhale, 2008) to provide a high probability of a positive treatment outcome. However, bacterial infections often require higher doses of the antibiotic to achieve therapeutic success *in vivo* (Kowalska-Krochmal and Dudek-Wicher, 2021). The effectiveness of the therapy depends on multiple factors for example: the type and concentration of antibiotic, exposure time to antibiotic, drug penetration to the site of infection and the duration of the infection before the drug treatment is delivered (Kowalska-Krochmal and Dudek-Wicher, 2021). As MIC is established on bacteria cultures *in vitro* against planktonic (free-living) bacteria, it does not consider these tissue-specific factors that affect the outcome of antibiotic treatment. Consequently, MIC antimicrobials are often found ineffective against persistent infections such as bacterial keratitis which involve biofilm formation (Costerton et al., 1987, Lebeaux et al., 2014, Davies, 2003). Treating biofilms often requires much higher than MIC of antibiotics, which can pose a risk of cytotoxicity. While some antibiotics are toxic to corneal epithelium (e.g. gentamicin), others can delay epithelial healing (ciprofloxacin), which can lead to corneal haze or keratolysis. Preservatives (e.g. benzalkonium chloride) in topical ophthalmic medications are directly cytotoxic to both host and pathogen cells but can improve antimicrobial efficacy by increasing drug penetration through devitalized epithelium (Eun et al., 1994, Noecker, 2001, Goldstein et al., 2022). To account for tissue-specific factors, therefore, a high throughput, *in vitro* model that can report on both the potency of the tested antibiotics and any tissue-specific response is needed to identify novel antimicrobials with suitable activities.

Currently, there is no ideal *in vitro* model for testing the efficacy of existing and new antimicrobial treatments. Keratitis models *in vivo* are not suitable for high throughput screening, are expensive, lead to animal suffering and therefore raise ethical concerns (Urwin et al., 2020). *Ex vivo* cornea infection models could be a good alternative to current *in vitro* techniques and have the potential to reduce and refine the use of animals for *in vivo* testing. However, *ex vivo* models are a relatively new concept and therefore our goal is to standardise and validate an *ex vivo* keratitis model for testing novel treatments. In this study, we used a previously established *ex vivo* porcine keratitis model (Okurowska et al., 2020) to test the activity of commonly used antibiotics. *P. aeruginosa* isolates PA14 and PA01 were selected because the biofilm formation (Wozniak et al., 2003, Colvin et al., 2011) and genetic similarities between these two strains (Lee et al., 2006) are well described in the literature. Each of these clinical isolates belongs to one of two major phylogenetic groups: group 1, which includes strain PA01, and group 2, which includes strain PA14 (Freschi et al., 2019). Each phylogenetic group is suspected to have a different effect on the host cells (Hilliam et al., 2020) and the clinical outcomes (Vallas et al., 1996, Borkar et al., 2013, Fleiszig et al., 1996, Lee et al., 2003a). Strain PA01 is moderately virulent and forms more structured biofilms on solid surfaces (Goodman et al., 2004) while PA14 is highly virulent, more cytotoxic and forms a weaker biofilm (Wiehlmann et al., 2007, Kasetty et al., 2021, Mikkelsen et al., 2011) called a pellicle that is associated with a stagnant liquid surface (Friedman and Kolter, 2004). Kasetty, S. et al. (2021) described differences in biofilm invasion strategies between these two strains in more detail. Genes encoding virulence factors in these strains are regulated by quorum-sensing (QS) systems, which are also well-described in the literature (Girard and Bloemberg, 2008, de Kievit, 2009). In this study, we wanted to determine if differences between these two strains will be obvious during different stages of *ex vivo* infection and after treatments with antibiotics. We demonstrate that our *ex vivo* porcine keratitis model can be used as a tool to test the effectiveness and optimal concentrations of new drugs or preservatives for ocular infections quickly, at a lower expense before these treatments are further validated *in vivo*. Our *ex vivo* model could help to select therapeutics that have a greater chance of success *in vivo*.

## MATERIALS AND METHODS

### Bacterial strain used

Two wild-type strains of *Pseudomonas aeruginosa* (PA01 and PA14) were a kind gift from Prof. Urs Jenal, University of Basel, Switzerland. Both strains were used to infect *ex vivo* porcine corneas and to establish MIC and MBEC values.

### MIC assay

The MIC value for *P. aeruginosa* PA01 and PA14 was determined according to the EUCAST guidelines (Hasselmann, 2000). The bacterial strains were inoculated in Mueller-Hinton cation-adjusted broth (MHB) for 24 hours at 37 °C with agitation at 110 rpm. Before each experiment 10 µl of 6-fold dilutions of the inoculum were spot-plated on blood agar plates, and the plates were incubated (Infors HT Multitron, UK) overnight at 37 °C to enumerate colony-forming units in the inoculum. Two hundred microliters of MHB containing an inoculum with 3x10^5^ CFU per well and different concentrations of the test antibiotics were added to each well in a 96-well plate. A concentration of antibiotics ranging from 0.006 to 32 µg/µL was tested. The MIC value was determined as the lowest concentration of an antibiotic which completely inhibits visible bacterial growth after 24 hours at 37 °C in static conditions. In total six antibiotics were tested: gentamicin, meropenem and ciprofloxacin. The optical density at 600 nm was measured using the TECAN Spark plate reader (TECAN, Switzerland) to confirm the growth inhibition. One column of each 96-well plate was designated for growth control and one for sterility control. The procedure was repeated three times across different days for each antibiotic.

### Biofilm susceptibility and MBEC assays

Biofilm susceptibility testing was performed using a Calgary device (Innovotech, Canada) where the biofilm was grown on a peg (Harrison et al., 2010). First, growth conditions were verified by an equivalence test for biofilm formation (Figure 1 in Supplementary Materials) as described by Harrison et al. (2010). The bacterial strains were streaked out on an LB agar plate from cryogenic stock and incubated overnight at 37 °C. A single colony from the agar subculture was used to inoculate 5 mL Mueller Hinton Broth (MHB) and the suspension was incubated in a 50 mL Falcon tube while shaking at 110 rpm for 24 hours at 37 °C (Infors HT Multitron, UK). The bacterial suspension was centrifuged at 4000 g in Eppendorf 5710R (Thermo Fisher, UK) for 5 minutes. After discarding the supernatant, the pellet was resuspended in 5 mL of sterile MHB. The inoculum size was prepared in a fresh centrifuge tube by diluting the suspension of bacteria to an optical density (OD) of 0.05 at 600 nm. The OD_600nm_ was measured using the spectrophotometer Jenway (VWR, UK). The inoculum was pipetted in a 96-well plate with a final concentration of 8x10^6^ CFU of *P. aeruginosa* PA01 or PA14 per well (150 µl inoculum in each well). One column in a 96-well plate was used as a control and contained media without bacteria added. Pegs from the Calgary Device were immersed in the inoculum. The 96-well plate was double sealed with parafilm, placed inside a plastic box to reduce evaporation and incubated (statically) overnight at 37^0^C with 70% humidity in the incubator (Infors HT Multitron, UK) to allow biofilm formation on pegs. Before each experiment 10 µl of 6-fold dilutions of the inoculum was spot-plated on blood agar plates, and the plates were incubated overnight at 37 °C in order to enumerate colony forming units (CFU) in the inoculum. After overnight incubation, the pegs were rinsed twice for 1 minute in two 96-well plates with 200 µl of sterile water per well to remove bacteria that did not attach to the pegs (planktonic cells).

**Figure 1.**
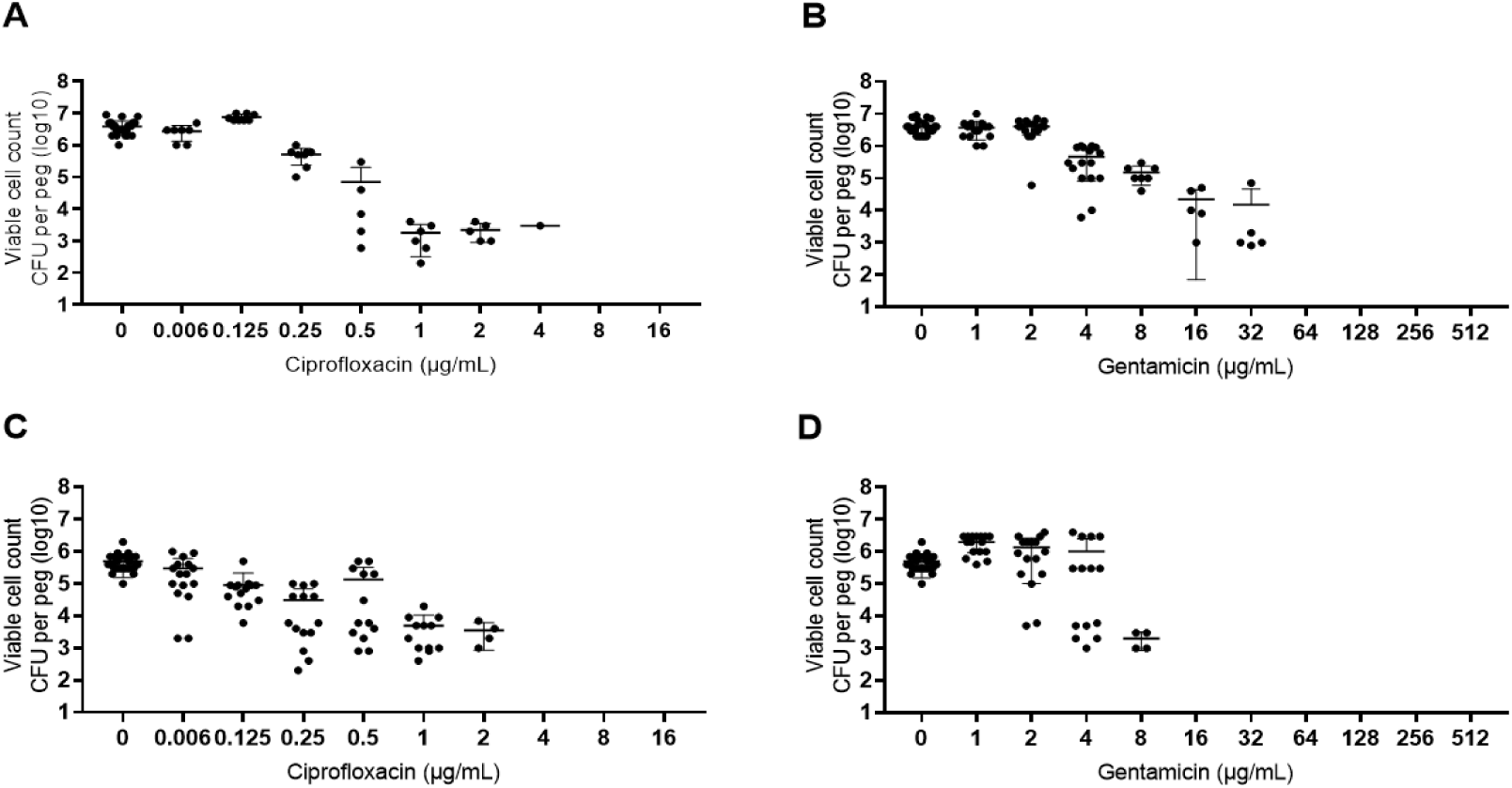
MBEC assay results representing colony forming units of P. aeruginosa PA01 (A, B) and PA14 (C,D) treated with ciprofloxacin and gentamicin.

For equivalence assay, the pegs were then transferred to a 96-well plate with 200 µl of LB with 1% Tween 20 per well, sonicated for 10 minutes at 60 Hz to disrupt bacteria from the biofilm on pegs into a recovery medium. After sonication, 20 µl of the recovery medium with the bacteria was diluted in series up to 10^4^ in 180 µl of sterile water. All dilutions were plated out on LB agar plates for CFU count and incubated at 37 °C overnight.

For the MBEC assay, the pegs were transferred after rinsing steps to a 96-well plate with antibiotics in MHB. The plate was incubated overnight and then rinsed and sonicated in the same way as equivalence assay plates. Ciprofloxacin, meropenem and gentamicin were tested with concentrations starting from 1µg µL^-1^ to 512 µg µL^-1^. The MBEC was determined as the wells with the lowest concentration of an antibiotic where the biofilm was completely eradicated i.e. there was no growth from biofilms across all replicates. One column of each 96-well plate was designated for untreated control and one for sterility control. The procedure was repeated four times across different days for each antibiotic with four technical replicates each time.

### Testing antibiotics on an *ex vivo* porcine cornea model

In this study, porcine eyes were extracted within four hours from slaughter and transported from the abattoir (R.B. Elliott and Son Abattoir, Calow, England) in a Nalgene container filled with sterile phosphate buffer saline (PBS, Sigma, Germany). The age of pigs varied between 26 to 28 weeks. The corneas were excised in the laboratory within two hours from delivery and used for experiments within a week from excision. The pigs were sacrificed for human consumption and not for the purpose of this study.

Porcine eyes were prepared for infection as described previously (Okurowska et al., 2020). Initially, the infection outcome at 1, 2, 4, 6 and 24 hours was tested to decide on the best time for addition of a treatment (Figure 3 in Supplementary Materials). The corneas were washed once with 2 mL PBS and incubated in an antimicrobial-free medium for 48 hours to remove residual antibiotics from the preparation procedure. The medium was replaced every day during this time. On the infection day, porcine corneas were infected with 8x10^6^ CFU in 200 µl of PBS and incubated for 6 hours. The inoculum size was established as described (Figure 2 in Supplementary Materials). After the incubation, the PBS along with the suspended bacteria were removed with a sterile 1 mL pipette tip and replaced either with 200 μL of PBS (control corneas) or with a PBS with added antibiotic (treated corneas). The corneas were treated with either 1024 μg mL^-1^ or with MIC concentration of ciprofloxacin, meropenem, and gentamicin for 18 hours at 37 °C. All corneas were photographed (Figures 4 & 5 in Supplementary Materials) with a Dino-lite Xcope camera (AnMo Electronics Corporation, Taiwan). Ninety cornea images were independently blind scored for opacity by five people using the following grading system: 0 – no haze, cornea clear; 1 – faint opacity or cloudiness visible; 2 – cornea looks swollen, white or hazy patch clearly visible. All data were plotted using GraphPad Prism version 8.4.1.

**Figure 4.**
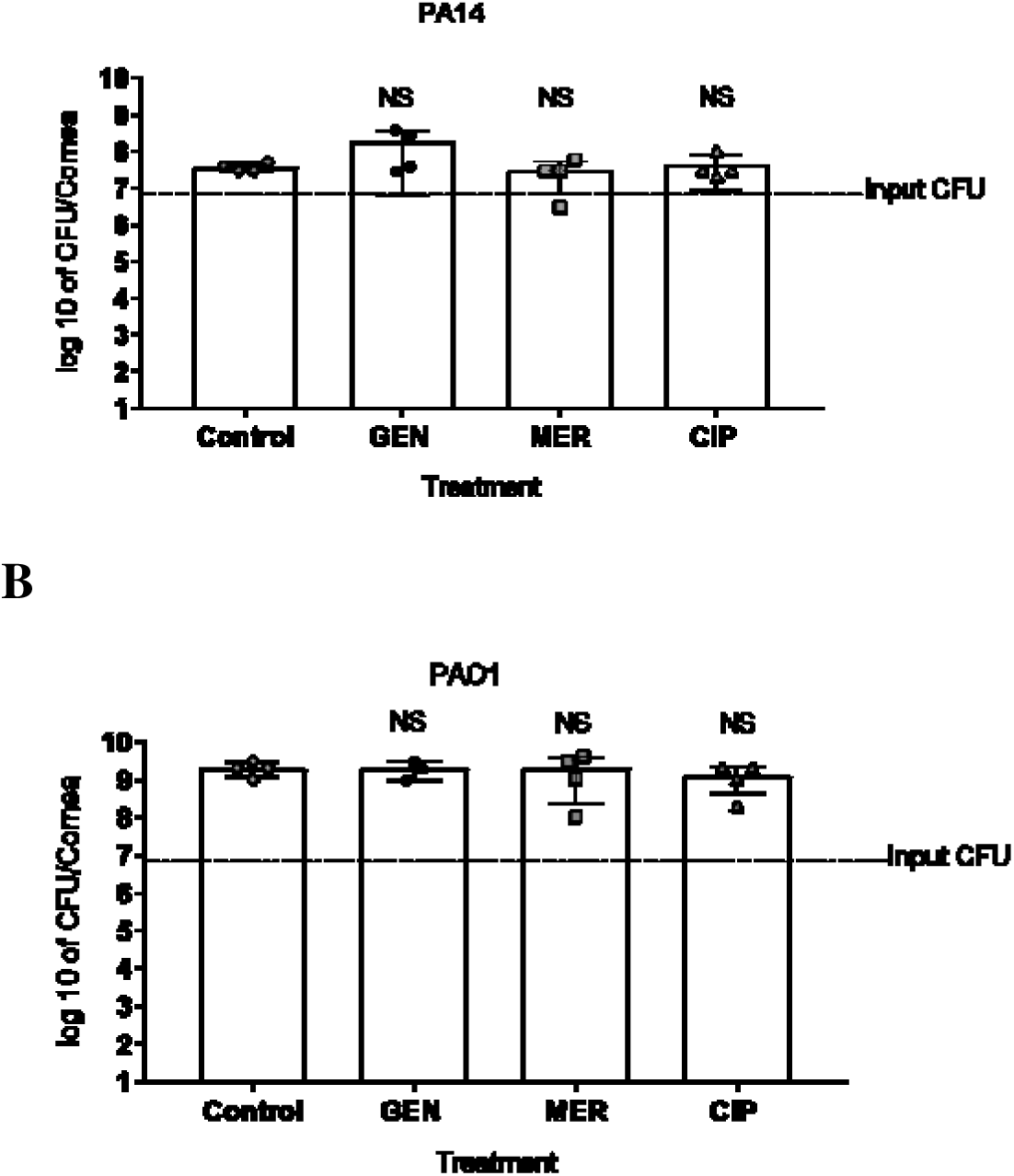
Number of CFU of *P. aeruginosa* in the *ex vivo* porcine corneas. Corneas were infected for 6 hours with (A) PA14 or (B) PA01. Control corneas were immersed in PBS while other corneas were treated with MIC concentrations of antibiotic dissolved in PBS. Following antibiotics were applied on infected corneas: gentamicin (GEN) (n = 4), meropenem (MER) (n = 4) and ciprofloxacin (CIP) (n = 4). Error bars are means ± SD. Kruskal-Wallis multiple comparison test was performed for the pairwise statistical analysis of treated against control colony forming units for each strain.

### Statistics

Statistical analysis of CFU between bacterial strains was carried out by unpaired t-tests with Holm-Sidak correction while effect of treatment versus placebo was calculated using Kruskal-Wallis multiple comparisons test, using GraphPad Prism version 8.4.1. *P*-values <0.05 were considered significant.

### Data availability

All supporting data are provided in the Supplementary Materials file.

## Results

### Antibiotic sensitivity in MIC assay

MIC assays revealed that both strains of *P. aeruginosa* used in this study were sensitive to gentamicin, meropenem and had intermediate resistance to ciprofloxacin (Table 1). The MIC values for gentamicin were identical for both strains (2 - 4 μg mL^-1^). However, some small differences were observed between strains treated with meropenem and ciprofloxacin. PA14 was marginally more susceptible to meropenem (0.25 μg mL^-1^) while PA01 was marginally more susceptible to ciprofloxacin (0.125-0.25 μg mL^-1^).

**Table 1.** Determination of MIC and MBC of *P. aeruginosa* for PA01 and PA14 isolates against gentamicin, meropenem and ciprofloxacin. Values in the table represent μg mL^-1^.

| Generic name<br>(class)<br>Break points<br>(EUCAST, 2022) | PA01 |  | PA14 |  | Mechanism of action |
| --- | --- | --- | --- | --- | --- |
|  | MIC | MBE<br>C | MIC | MBE<br>C |  |
| Gentamicin<br>(aminoglycoside)<br>$\leq 4$ S; $\geq 16$ (R) | 2-4 | 64<br>(16X -<br>32X<br>MIC) | 2-4 | 16<br>(4X -<br>8X<br>MIC) | Broad spectrum, inhibits<br>synthesis of bacterial proteins by<br>binding to 30S ribosomes |
| Meropenem<br>(carbapenem)<br>$\leq 2$ (S); $\geq 8$ (R) | 0.5-1 | >512 | 0.25 | >512 | Broad spectrum, inhibition of<br>bacterial cell wall synthesis |
| Ciprofloxacin<br>(fluoroquinolone)<br>$\leq 0.001$ (S); $\geq 0.5$<br>(R) | 0.125-0.25 | 4-8<br>(16X -<br>64X<br>MIC) | 0.25-0.5 | 4<br>(8X -<br>16X<br>MIC) | Inhibits DNA replication by<br>inhibiting bacterial DNA<br>topoisomerase and DNA-gyrase |

### Antibiotic sensitivity in MBEC assay

Despite MIC results showing sensitivity to meropenem (Table 1), biofilms of both *P. aeruginosa* strains demonstrated tolerance to meropenem exceeding the concentrations tested (>512 μg mL^-1^). For the PA01 strain, MBEC values were 16-64 times higher than MIC for ciprofloxacin and 16-32 times higher than MIC for gentamicin (Figs. 1A and 1B). MBEC values for PA14 strain for ciprofloxacin were 8-16 times higher than MIC and 4-8 times higher than MIC for gentamicin (Fig. 1C and 1D). These results suggest that the biofilm on pegs formed by strain PA14 was more sensitive to gentamicin and ciprofloxacin compared to strain PA01 (Table 1). MBEC testing made the difference between the two strains more noticeable. With reference to the breakpoint system (Table 1) and subsequent clinical relevance, the MBEC results demonstrate that biofilms formed by *P. aeruginosa* could be classified as resistant to gentamicin, meropenem and ciprofloxacin.

### Investigation of antimicrobial efficacy on the *ex vivo* porcine keratitis model

#### Testing antibiotics on the *ex vivo* porcine keratitis model

Firstly, the effect of MIC concentrations of antibiotics on infected tissue was investigated. *Ex vivo* porcine corneas were infected on average with 1 x 10^7^ CFU *P. aeruginosa* PA14 and 9 x 10^6^ *P. aeruginosa* PA01 for 6 hours and then MIC concentrations of gentamicin, meropenem and ciprofloxacin were applied for 18 hours. While MIC concentrations of antibiotics successfully inhibited the growth of bacteria *in vitro*, these concentrations were ineffective (p >0.05) for both tested strains of *P. aeruginosa* PA14 and PA01 in the *ex vivo* porcine cornea model (Fig. 4). Raw data are presented in Supplementary Materials Table 1. This demonstrates that the application of MIC concentrations of these antibiotics on *ex vivo* cornea is insufficient to treat ocular infections with *P. aeruginosa* even though the infected tissue was continually exposed to the antibiotic for 18 hours.

The concentration of antibiotics (gentamicin, meropenem and ciprofloxacin) that were applied on *ex vivo* porcine corneas was increased to 1025 μg mL^-1^. This concentration is 256 times MIC for gentamicin for strains PA01 and PA14, respectively. For meropenem, this concentration is 1025 times MIC for strain PA01 and 4100 times MIC meropenem for PA14. For ciprofloxacin, this concentration is 4100 times MIC for strain PA01 and 2050 times MIC for strain PA14. As this concentration is higher than MIC (and MBEC) some growth inhibition on *ex vivo* infected tissue was expected. A significant reduction in bacteria load for strain PA01 in corneas treated with gentamicin (p = 0.0051), meropenem (p < 0.0001) and ciprofloxacin (p < 0.0001) was observed when compared to controls (Fig. 5). In contrast, there was no significant reduction for corneas infected with strain PA14 and treated with gentamicin (p = 0.15) whereas, treatment of PA14 with meropenem (p = 0.0001) and ciprofloxacin (p <0.0001) had a noticeable reduction in bacteria load (Table 2 in Supplementary Materials).

**Figure 5.**
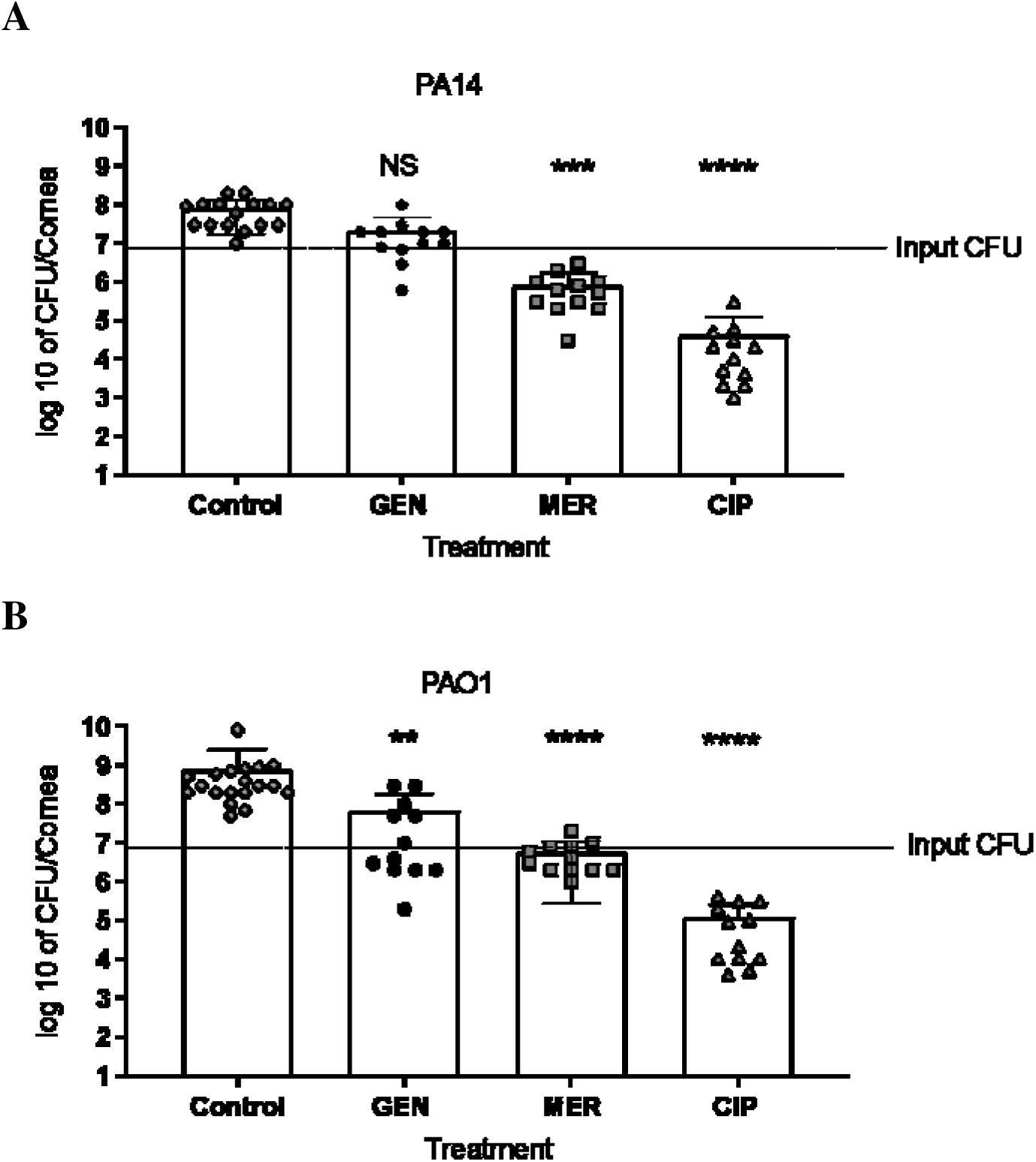
Colony forming units of *P. aeruginosa* in the *ex vivo* porcine corneas infected for 6 hours with (A) PA14 or (B) PA01. Control corneas (n = 19 & 16) were immersed in PBS while other corneas were treated with 1025 μg mL^-1^of antibiotic dissolved in PBS. Following antibiotics were applied on infected corneas: gentamicin (GEN) (n = 12), meropenem (MER) (n = 12) and ciprofloxacin (CIP) (n = 12). Error bars are means ± SD. Kruskal-Wallis multiple comparison test was performed for the pairwise statistical analysis of treated against untreated colony forming units for each strain; significant difference (*p* value < 0.05) is denoted with *.

#### Assessing opacity

The presence and intensity of discolouration and opacity for ninety-nine corneas were undertaken by assessing the images (Supplementary Materials Fig. 4 & 5) to verify if *infection* or the effect of treatment could be determined visually. The images of corneas were allocated to the following three grades: 0 – the corneas looked clear and not infected; 1 – corneas looked infected, slightly hazy and cloudy; 2 – corneas looked infected, swollen and white/milky in colour. Images of corneas were blind-scored by five different people and presented as percentages (Fig. 6).

**Figure 6.**
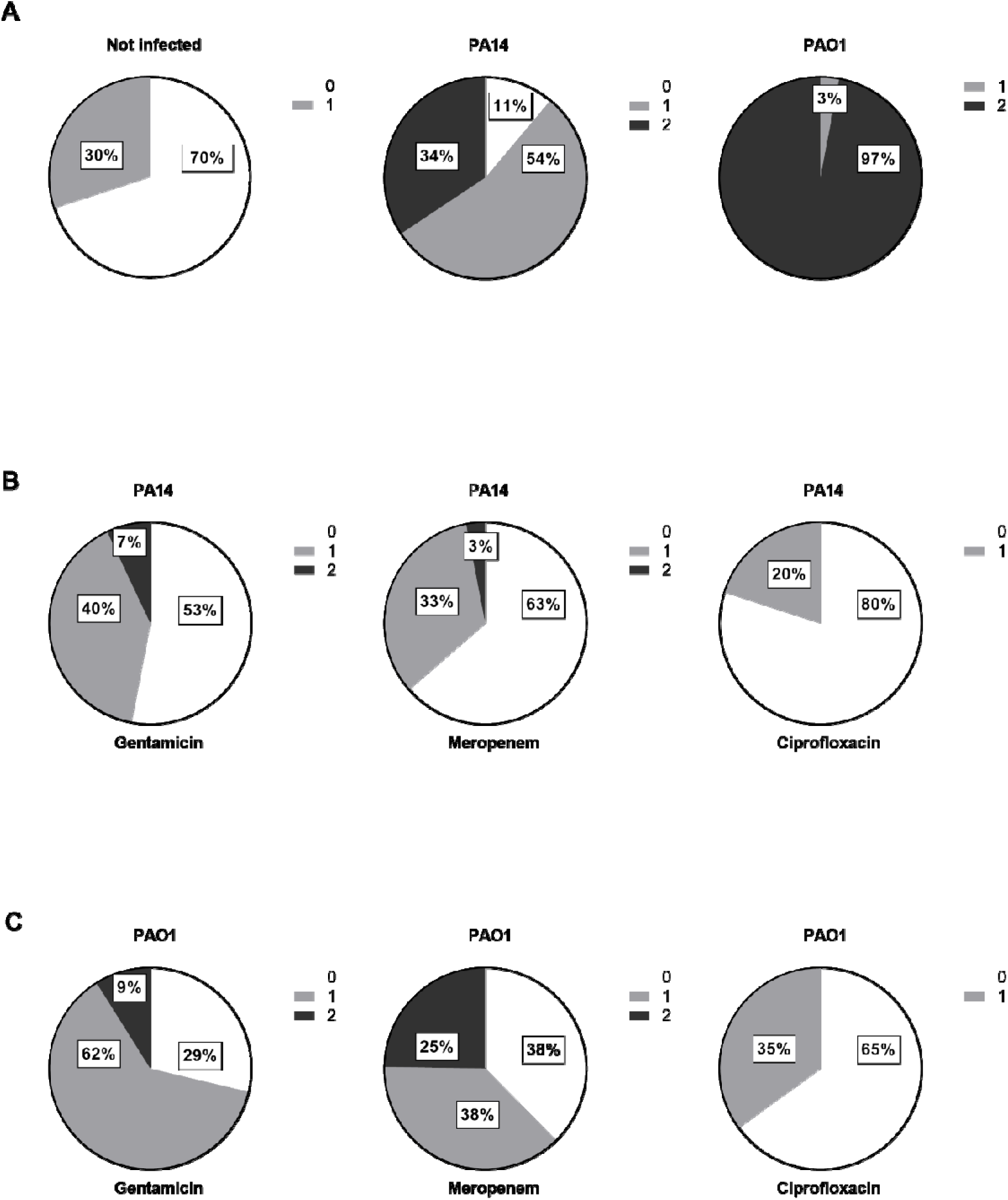
Effects of antibiotic treatment on infected cornea. The percentage of scored *ex vivo* porcine corneas (n = 99) that were not infected (n = 4) versus infected with *P. aeruginosa* strain PA14 (n = 42) and PA01 (n = 44) for 24 hours (A). Some corneas were treated with 1025 μg mL^-1^ of gentamicin, meropenem or ciprofloxacin after infection with PA14 (B) and PA01 (C). Corneas were allocated to the following groups: 0 – corneas clear, no haze or infection visible; 1- corneas looked infected, slightly hazy and cloudy; 2 – corneas looked infected, swollen and white/milky in colour.

Most uninfected corneas (70%) were correctly identified while 30% were graded 1 (Fig. 6A). This is because the scoring of uninfected corneas could have been affected by swelling that naturally occurs during incubation in media and could make some corneas appear less transparent than normal. Lack of previous experience in scoring, changes in the natural light in the room and quality of some images could also influence cornea grading.

Some untreated corneas infected with strain PA14 looked clear (11%) whilst the majority were correctly identified as infected (54% scored grade 1 and 34% scored grade 2) (Fig. 6A). Majority of untreated corneas (97%) were correctly identified as being infected with strain PA01 due to the development of an obvious haziness (grade 2) suggesting infection with strain PA01 results in ulceration and severe tissue damage on *ex vivo* porcine cornea (Fig. 6A).

Overall, corneas infected with strain PA14 and treated with antibiotics (Fig. 6B) developed less hazy in comparison to strain PA01 (Fig. 6C). Smaller percentage of cornea infected with either strain and treated with 1025 µg mL^-1^ gentamicin and meropenem were scored grade 2 in comparison to untreated (Figs. 6B & 6C versus Fig. 6A). None of the corneas infected with either strain and treated with ciprofloxacin was graded 2 and the majority were graded 0 which suggests that ciprofloxacin had a beneficial effect on preserving corneal transparency.

Opacity scoring demonstrated visual differences in opacity between infection with *P. aeruginosa* cytotoxic strain PA14 and infection with invasive strain PA01. The PA01 strain showed the most visually obvious reduction in opacity for both untreated as well as treated corneas. However, higher transparency of corneas treated with antibiotics should not be solely used to determine the effectiveness of a drug because, for example, in the case of gentamicin the reduction in colony-forming units was very minimal even though the transparency was improved. These data demonstrate that images of *ex vivo* corneas could be used as one of the preliminary indicators showing the immediate response of infection to a treatment.

## Discussion

We previously established an *ex vivo* porcine model of *Pseudomonas aeruginosa* keratitis (Okurowska et al., 2020). In this study, we demonstrate that our *ex vivo* porcine cornea model can be used for testing novel treatments against keratitis. We first established MIC and MBEC values for gentamicin, meropenem and ciprofloxacin using cytotoxic (strain PA14) and invasive (strain PA01) strains of *P. aeruginosa*. Next, we tested the input of bacteria needed to develop an infection and then monitored the development of an infection over time. Finally, we investigated differences in response to antibiotic treatments between cytotoxic (strain PA14) and invasive (strain PA01) strains of *P. aeruginosa* on the *ex vivo* porcine keratitis model.

Comparing MIC and MBEC data to literature was challenging because of the variance in experimental protocols between research groups (Kowalska-Krochmal and Dudek-Wicher, 2021, Schuurmans et al., 2009) but overall, our findings followed the trend in literature. The MBEC values of all tested antibiotics were much higher in comparison to MIC. Our results tie in well with previous studies showing that higher concentrations of antibiotics are needed to eradicate biofilms compared to their planktonic counterparts (Bagge et al., 2004, Brady et al., 2017, Bowler et al., 2012, Ceri et al., 1999).

Gentamicin is used in the early stages of keratitis (Dart and Seal, 1988). We found that MIC results in our study were in alignment with those found in the literature and demonstrated that both strains of *P. aeruginosa* (PA14 and PA01) were equally sensitive to gentamicin. While MIC values did not demonstrate any obvious differences between the investigated strains, MBEC results clearly show that PA01 was more resistant to gentamicin in comparison to PA14. Like other studies, MBEC values showed that biofilms were resistant to gentamicin in *P. aeruginosa* (Bowler et al., 2012, Ceri et al., 1999, Billings et al., 2013). These results suggest that whilst gentamicin could be used to eradicate planktonic forms of *P. aeruginosa*, treating the biofilm formed by this bacterium would require much higher concentrations.

Meropenem has good corneal penetration and low cytotoxicity (Sueke et al., 2015), with promising results in an *in vivo* rabbit keratitis model. Our MIC values for *P. aeruginosa* PA01 showed sensitivity towards meropenem and were identical (0.5 μg mL^-1^) (Riera et al., 2010, Ocampo-Sosa et al., 2012) or close (1-2 μg mL^-1^) (Haagensen et al., 2017, Bowler et al., 2012, Monahan et al., 2014) to those found in the literature. The MIC values for strain PA14 in our study are also in accordance with findings reported by others (0.25 μg mL^-1^)

(Hassan et al., 2020, Ocampo-Sosa et al., 2012). MBEC values in literature for *P. aeruginosa* PA01 biofilm treated with meropenem were consistently much higher than MIC values (Haagensen et al., 2017, Bowler et al., 2012). Thus, our MBEC results agree with previous studies showing resistance in *P. aeruginosa* towards this antibiotic (Bowler et al., 2012, Haagensen et al., 2017). These findings support the notion that meropenem is more effective against actively dividing, planktonic bacteria or early-stage biofilm, while less effective against established biofilms (Bowler et al., 2012).

Finally, we investigated ciprofloxacin in our studies because it is considered one of the most effective antibiotics against *P. aeruginosa* and therefore used as a first-line treatment in the UK (Hilliam et al., 2020). Our study demonstrated that ciprofloxacin was certainly the most potent antibiotic against planktonic and biofilm of *P. aeruginosa* not only *in vitro* but also in *ex vivo* cornea models. Our MIC values for ciprofloxacin indicate that both strains were susceptible, with PA14 marginally more resistant than PA01. Overall, the results for MIC were similar to those in the literature (Riera et al., 2010) (Shafiei et al., 2014, Bowler et al., 2012) ^(^Fernandez-Olmos et al., 2012) for PA01 and PA14. (Soares et al., 2019, Bruchmann et al., 2013). Our MBEC results also match trends in other studies *in vitro* which demonstrated that there was a small but not significant difference in response to ciprofloxacin between biofilm formed by PA14 and PA01 (Billings et al., 2013, Benthall et al., 2015).

In initial studies to establish an *ex vivo* keratitis model, we tested various bacterial loads in the inoculum to initiate infection. It was found that as few as 215 CFU of *P. aeruginosa* PA14 per cornea was enough to initiate infection in our *ex vivo* model and the CFU retrieved from the cornea after 24 hours were the same regardless of the input. We speculate that this phenomenon happens at 24 hours post infection because the nutrients become more limited at this time point and similarly to batch cultures, the bacteria reached a stationary growth phase (Llorens et al., 2010, Rolfe et al., 2012). Lower inoculum was found to decrease the prevalence of ulcerative microbial keratitis in live animals (Lawinbrussel et al., 1993) therefore researchers often initiate infection using an inoculum load higher than or equal to 1 x 10^6^ CFU of *Pseudomonas* sp. per eye (Tam et al., 2011, Lawinbrussel et al., 1993, Preston et al., 1995, Augustin et al., 2011). Additionally, as previously discussed (Urwin et al., 2020), we are unable to compare our results to other studies *ex vivo* due to lack of standardised protocol. As a higher inoculum allows reliable bacterial quantification and makes bacterial visualisation on infected corneas easier (Ting et al., 2021a) we subsequently used an inoculum containing at least 1 x 10^6^ CFU per cornea in further experiments.

To identify whether it is possible to distinguish between infections caused by cytotoxic and invasive strains of *P. aeruginosa*, we compared progress of infection over time between cytotoxic *P. aeruginosa* PA14 and invasive *P. aeruginosa* PA01 strain by monitoring the number of CFU retrieved from the cornea over time. Growth plateaux observed after 18 hours of incubation with both strains suggests that stationary phase was reached for both strains at this point. This suggests that the enhanced cytotoxicity of *P. aeruginosa* PA14 did not seem to confer a selective advantage during infection of the wounded *ex vivo* porcine cornea. These observations led us to conclude that enhanced cytotoxicity did not dramatically affect progress of infection in our porcine keratitis model.

Finally, when the antibiotics were tested on the *ex vivo* keratitis model, we discovered that gentamicin was not effective at a concentration of 1.25 mg mL^-1^. Studies on rabbits *in vivo* used various concentrations of gentamicin (1.6, 3, 5 &13 mg mL^-1^) to treat *P. aeruginos*a keratitis with mixed therapeutic outcomes (Punitan et al., 2019, Rootman and Krajden, 1993, Fruchtpery et al., 1995, Silbiger and Stern, 1992, Gupta et al., 1995, Kowalski et al., 2013, Ahmad et al., 1977). Some of the concentrations used in the mentioned studies were cytotoxic because it was found that as little as 3 mg mL^-1^ of this antibiotic is cytotoxic in human corneal epithelial cells in vitro (Tsai et al., 2010) and impairs the wound healing in rabbits in vivo (Stiebel-Kalish et al., 1998). This suggests that the potential harm of higher gentamicin concentrations used to treat infections may outweigh the benefit, especially with prolonged exposure.

Meropenem has low toxicity, and good corneal tissue penetration (Sueke et al., 2015) and it was found to be very effective in *Pseudomonas* keratitis in concentrations of 50 mg mL^-1^ in rabbits *in vivo* (Bozkurt et al., 2021) and humans (Pande and Bhailume, 2014) without any side effects. Some studies found that meropenem concentrations of 5 mg mL^-1^ increased cellular activity in corneal epithelial cell lines and the cell viability was still high (96%) after meropenem treatment (Sueke et al., 2014, Sueke et al., 2015). Meropenem reduced the bacterial load in our *ex vivo* corneas similarly to other studies on *ex vivo* rabbits or humans (Bozkurt et al., 2021). Low toxicity at high concentrations and reduction of bacteria load in studies *in vivo* suggest that meropenem could be a good alternative drug against keratitis in the future (Pande and Bhailume, 2014, Bozkurt et al., 2021). However, the resistance towards this antibiotic in MBEC data is concerning (Bowler et al., 2012, Haagensen et al., 2017). Haagensen et al. (2017) demonstrated that meropenem was highly effective in the early stages of *P. aeruginosa* PA01 biofilm formation. We also achieved a good reduction of bacteria load after application of meropenem after 6 hours post-infection, during possibly early stages of biofilm formation. More studies need to be conducted to assess the effectiveness of this drug and the possibility of its use in clinical practice. Some studies report that combining meropenem and ciprofloxacin can have a synergistic effect against some clinical isolates of *P. aeruginosa* (Erdem et al., 2002, Erdem et al., 2003, Siqueira et al., 2014, Pankuch et al., 2008, GarciaRodriguez et al., 1996) which could be tested on our *ex vivo* porcine keratitis model in the future.

Ciprofloxacin has a very good tissue penetration property. Exposure to this antibiotic for as short as 10 minutes has been demonstrated to result in concentrations exceeding MIC in human cornea *ex vivo* (Silva et al., 2017, McDermott et al., 1993, Akkan et al., 1997, Ozturk et al., 1999) therefore we suspect that the 18-hour continuous exposure to this antibiotic in our study very likely resulted in MIC concentration in corneal tissue. Ciprofloxacin was very effective in eradicating *P. aeruginosa* at higher concentrations in our experiments, which is in line with studies *in vivo*. The treatment was equally effective against cytotoxic and invasive strains of *P. aeruginosa*. Several studies found that treating corneas with ciprofloxacin significantly reduced or completely ceased infection with *P. aeruginosa* in live rabbits (Obrien et al., 1988, Aliprandis et al., 2005, Guzek et al., 1994, LaBorwit et al., 2001, Bu et al., 2007, Kowalski et al., 2001, Lauffenburger and Cohen, 1993, Oguz et al., 2005, Rhee et al., 2004) and humans (Levey et al., 1998). However it was found that phenotypic adaptation towards persistence to this antibiotic happens very early if supra-MIC concentrations are used and as a result, ciprofloxacin may fail to eradicate biofilm (Soares et al., 2019). Using higher concentrations of ciprofloxacin (0.3%) can cause crystalline corneal precipitation in humans (Wilhelmus and Abshire, 2003, Wilhelmus et al., 1993, McDonald et al., 2014).

Visual acuity is clinically one of the parameters showing therapeutic response in patients (Borkar et al., 2013, Hue et al., 2009). We observed that corneal damage caused by *P. aeruginosa* in our *ex vivo* keratitis model looked visibly similar to images found in clinical case reports (Hue et al., 2009). The response to different treatments can be observed and used to foresee the outcome which makes this model even more advantageous in comparison to other *in vitro* models. Genetic differences between cytotoxic and invasive *P. aeruginosa* strains led to different effects on epithelial cells (Fleiszig et al., 1997) which may be observed visually (Cole et al., 1998). However, other researchers found a lack of correlation between the number of viable bacteria remaining after antibiotic treatment and disease severity assessed visually from images (Lee et al., 2003b) in a similar way to ours. This was verified in our study where the corneas looked healthy after gentamicin treatment despite a high colony count. In our experiments, the invasive strain PA01 had initially the highest observable opacity with and without an antibiotic treatment in comparison to the cytotoxic strain PA14. A similar conclusion was reached by Borkar et al. (2013) where the ulcer size from invasive strains of *P. aeruginosa* in human keratitis was significantly bigger than from cytotoxic although genotypically invasive strains were associated with better visual acuity at enrolment. Some studies on mice showed that the damage in the centre of the cornea is not only due to bacterial damage but also a result of neutrophil infiltration (Fleiszig et al., 1996, Cole et al., 1998, Borkar et al., 2013) however our model *ex vivo* lacks neutrophils therefore ulceration comes from bacterial action.

The limitation of the present study naturally includes absence of fully operating host-defences in the *ex vivo* model. Nevertheless, the response to treatment with tested antibiotics was in line with trends found in literature and showed that observations of our *ex vivo* keratitis model is very similar to other animal models *in vivo* and to findings in clinical studies on humans. Therefore, our *ex vivo* porcine cornea model is a practical tool for rapidly and cost effectively screening the efficacy of ocular drugs with good sensitivity and reliability. We contend that our *ex vivo* model could be used to reduce and refine use of live animals in keratitis studies. The observations from our *ex vivo* keratitis model could advance discovery of new ocular drugs, facilitate their rapid translation to the market and serve as a guidance for clinical application in the future.

## Acknowledgements

We would like to thank R. B. Elliot and Son Abattoir for providing porcine corneas for research. The authors also want to thank Mr Jonathan Emery, Dr Mahendra Raut, Prof. Annette Taylor and Ms Hannah Regan for scoring cornea opacity. The authors would like to thank Dr Grace Crowther for her technical support.

## Author contributions

KO and EK conceived the hypothesis and research design, KO performed all the experiments, analysed the data, and wrote the manuscript. EK and PM provided resources and reviewed the manuscript.

## Ethical statement

The authors are accountable for all aspects of the work in ensuring the questions related to the accuracy and integrity of any part of the work are appropriately investigated and resolved. Pig eyes were obtained from animals sacrificed for human consumption and not for this study; therefore, ethical approval was not required.

