## Supplementary Materials for "Increased tolerance to commonly used antibiotics in a *Pseudomonas aeruginosa ex vivo* porcine keratitis model"

**Equivalence assay**

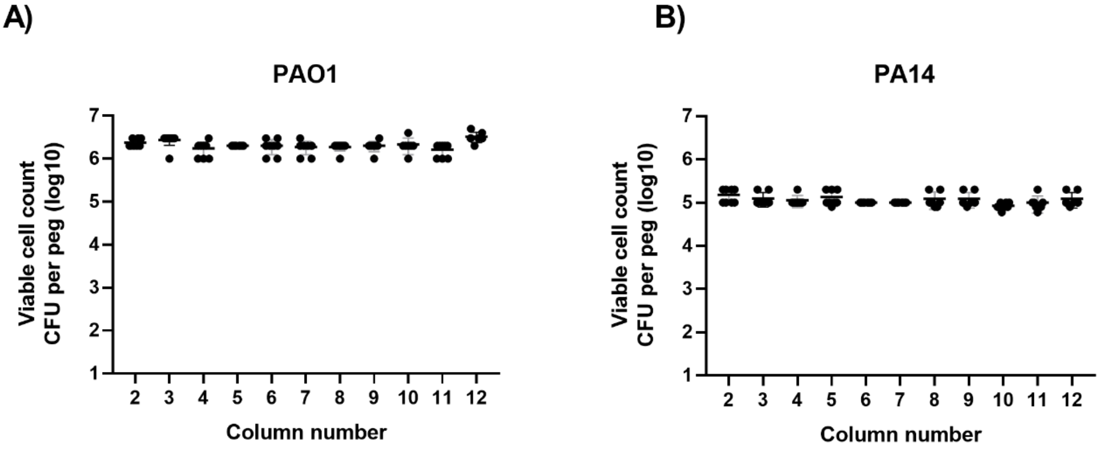


Figure 1. Equivalence assay results representing colony-forming units of *P. aeruginosa* PA01 (A) and PA14 (C) retrieved from pegs across all columns in a 96-well plate.

**Effect of inoculum size on final bacterial load**

To establish the inoculum size needed to initiate an infection in the porcine cornea, various CFUs of *P. aeruginosa* PA14 were added to wounded corneas. A viable count of bacteria retrieved from the infected cornea after 24 hours of infection (Fig. 2A) and 48 hours of infection (Fig. 2B) was carried out. Despite the starting inoculum size, an average of 6 x $10^8$ CFUs per cornea were retrieved after 24 hours and 2 x $10^9$ CFUs per cornea after 48 hours. There was no significant difference in CFU between groups and two incubation times. These results indicate that the ultimate bacterial load in the porcine *ex vivo* cornea infection model is independent of the initial bacterial load. Due to the good reproducibility in the number of CFU retrieved after infection with a higher starting inoculum size, in further experiments, an inoculum size of greater than 1 x $10^6$ CFU per cornea was aimed for. We established that the maximum incubation time for all following experiments was 24 hours because 48 hours of incubation resulted in complete lysis of the cornea by the bacteria.

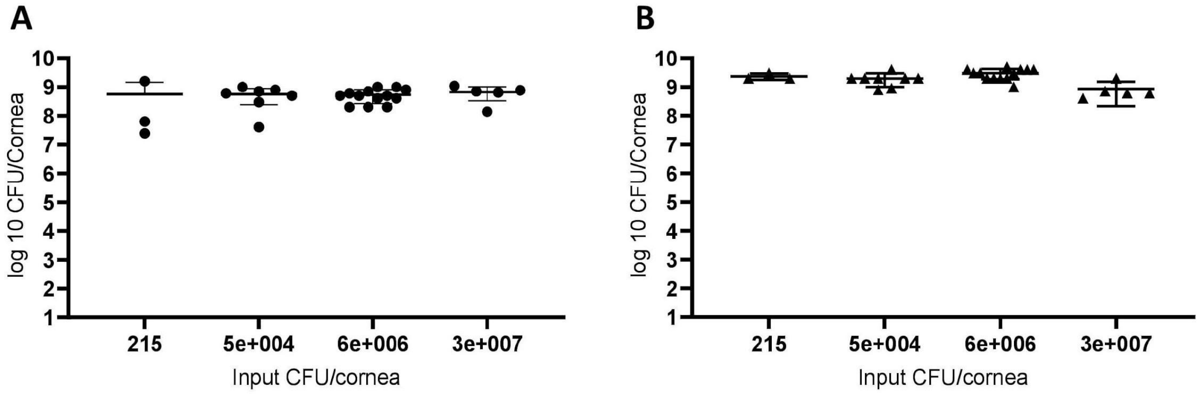


Fig 2. Number of viable *P. aeruginosa* PA14 retrieved from porcine cornea after infection with 215, 5x$10^4$, 6x$10^6$ and 3x$10^7$ CFU per cornea. Corneas were infected for 24 hours (A) and 48 hours (B). Each dot represents the results from one cornea. Error bars represent standard deviation. Statistical significance was calculated with the Kruskall-Wallis test followed by Dunn's multiple comparisons test *$p$-value <0.05.

**Effect of incubation time on the progress of infection**

To investigate the progress of infection over time, porcine corneas were infected with *P. aeruginosa* PA14 and *P. aeruginosa* PA01 and a viable count was carried out on bacteria retrieved from the infected cornea after 1, 2, 4, 6, 18 and 24 hours post-infection (hpi) (Fig. 3). With *P. aeruginosa* PA14, an average of 1.9 x $10^6$ CFU per cornea were retrieved after 1 hpi (n =7), 2.9 x $10^6$ CFU per cornea were retrieved after 2 hpi (n = 6), and 4.9 x $10^6$ CFU per cornea were retrieved after 4 hpi (n = 6) (Fig. 3A). At all these time points, the number of CFU retrieved per cornea were lower than the inoculum size (7.7 x $10^6$ CFU per cornea) reflecting the impact of post-incubation rinsing steps included in the protocol during which the bacterial population not securely adhered to the corneal tissue are removed. After 6 hpi, the number of bacteria retrieved from the infected cornea were approximately equal to the inoculum size despite rinsing (n = 6). Incubation beyond 6 hpi reproducibly resulted in a clear increase of CFU retrieved per cornea despite rinsing, resulting in 1.0 x $10^8$ CFU per cornea at 18 hpi (n = 6) and 9.0 x 10 $^7$ at 24 hpi (n = 6) (Fig. 3A). Difference in CFU values for PA14 retrieved at 1 hpi and 2 hpi in comparison to 18 hpi and 24 hpi was significant ($p<0.05$).

A similar trend was seen in the progress of infection in the *ex vivo* porcine cornea infected with *P. aeruginosa* PA01 strain (Fig. 3B). An average of 3.4 x$10^6$ CFU per cornea were retrieved after 1 hpi (n = 4), 2.2 x $10^6$ CFU per cornea were retrieved after 2 hpi (n = 14) and 4.1 x $10^6$ CFU per cornea were retrieved at 4 hpi (n= 6). Like the infection with *P. aeruginosa* PA14, at all these time points, the number of CFU retrieved per cornea were lower than the inoculum size (7.9 x $10^6$ CFU per cornea). Subsequently, the increase in bacteria load in the infected cornea was higher compared to the inoculum size for *P. aeruginosa* PA01 (Fig. 3C): 2.0 x $10^7$ CFU per cornea at 6 hpi (n = 6), 1.6 x $10^8$ CFU per cornea at 18 hpi (n = 4) and 1.7 x $10^8$ CFU per cornea at 24 hpi (n = 25) (Fig. 3B). Difference in CFU values for PA01 retrieved at 1hpi, 2 hpi and 4hpi in comparison to 18 hpi and 24 hpi was significant (p<0.05).

These data demonstrate that both strains of *P. aeruginosa* were able to initiate and maintain infection on porcine corneas within the first few hours of incubation. In both strains, despite the inclusion of a washing step, there was a net increase in the number of CFU retrieved after incubation compared to the inoculum which suggests that infection was well established in the model. In the subsequent experiments, antibiotic treatments were added to corneas at 6 hpi because there was a clearly visible increase in CFU counts at this time point in comparison to the input of bacteria which indicated that the infection was well-established.

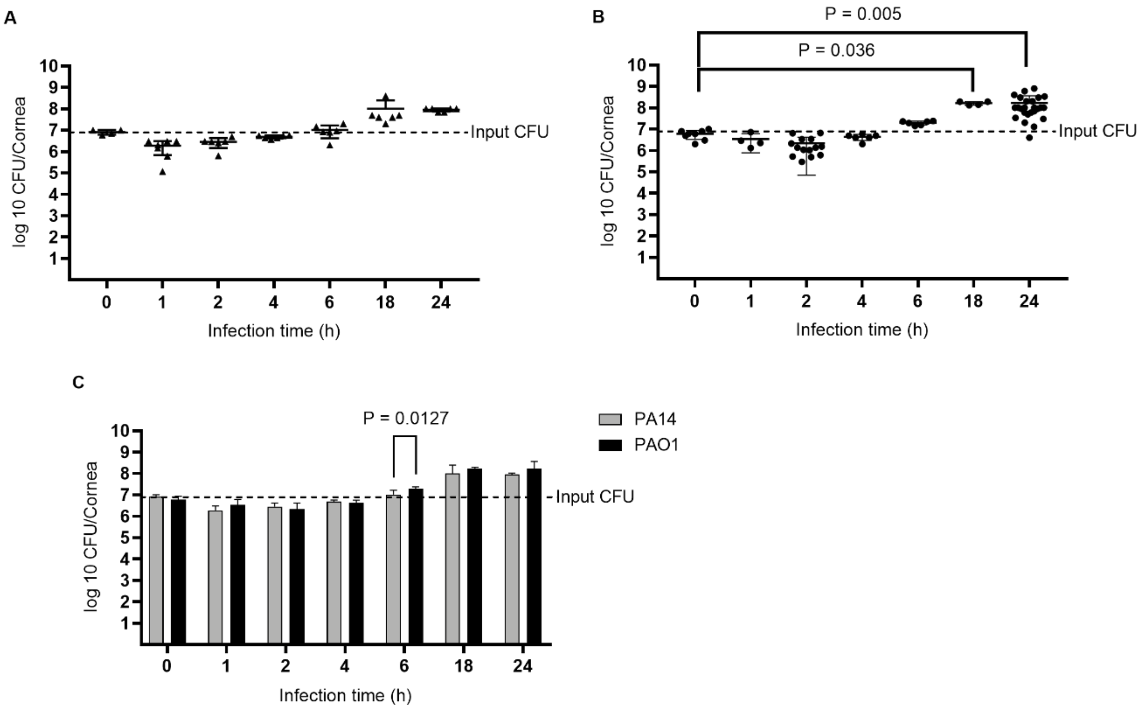


Fig. 3. Number of CFU of *P. aeruginosa* on porcine corneas. Corneas were infected for 1, 2, 4, 6, 18 and 24 hours with *P. aeruginosa* PA14 (A) and *P. aeruginosa* PA01 (B). Data from both strains are compared on one graph (C). Inoculum CFU are shown both as CFU at t=0 infection time (x-axis) as well as a dotted line labelled as Input CFU. Each dot on charts A and B represents one cornea. Error bars represent standard deviation. Statistical significance for graphs A and B is presented according to the Kruskall-Wallis test while for graph C is presented according to the unpaired *t*-test with Holm-Sidak correction **p*-value <0.05.

### Investigation of antimicrobial efficacy on the *ex vivo* porcine keratitis model

#### Testing antibiotics on the *ex vivo* porcine keratitis model

Table 1. Data summary of average colony forming units of *P. aeruginosa* in the ex vivo porcine corneas infected for 6 hours with PA01 or PA14 and treated with MIC concentrations of gentamicin, ciprofloxacin and meropenem.

**PA01**

| Treatment | Concentration $\mu g\ mL^{-1}$ | Average | N | SD |
|---|---|---|---|---|
| PBS | 0 | 3.E+09 | 4 | 8.E+08 |
| Gentamicin | 4 | 2.E+09 | 4 | 8.E+08 |
| Ciprofloxacin | 0.25 | 2.E+09 | 4 | 1.E+09 |
| Meropenem | 1 | 1.E+09 | 4 | 9.E+08 |

**PA14**

| Treatment | Concentration $\mu g\ mL^{-1}$ | Average | N | SD |
|---|---|---|---|---|
| PBS | 0 | 4.E+07 | 4 | 1.E+07 |
| Gentamicin | 4 | 2.E+08 | 4 | 2.E+08 |
| Ciprofloxacin | 0.5 | 3.E+07 | 4 | 2.E+07 |
| Meropenem | 0.25 | 5.E+07 | 4 | 5.E+07 |

Table 2. Data summary of average colony forming units of *P. aeruginosa* in the *ex vivo* porcine corneas infected for 6 hours with PA01 or PA14 and treated with gentamicin, ciprofloxacin and meropenem.

**PA01**

| Treatment | Concentration mg/mL | Average CFU | SD | N | % Reduction | LOG Reduction |
|---|---|---|---|---|---|---|
| PBS | 0 | 8.E+08 | 2.E+09 | 19 | | |
| Gentamicin | 1.025 | 7.E+07 | 1.E+08 | 12 | 91.260 | 1 log |
| Ciprofloxacin | 1.025 | 1.E+05 | 1.E+05 | 12 | 99.985 | 5 log |
| Meropenem | 1.025 | 5.E+06 | 4.E+06 | 12 | 99.340 | 2 log |

**PA14**

| Treatment | Concentration mg/mL | Average CFU | SD | N | % Reduction | LOG Reduction |
|---|---|---|---|---|---|---|
| PBS | 0 | 1.E+08 | 6.E+07 | 16 | | |
| Gentamicin | 1.025 | 2.E+07 | 2.E+07 | 12 | 79.266 | < 1 log |
| Ciprofloxacin | 1.025 | 5.E+04 | 2.E+04 | 12 | 99.955 | 4 log |
| Meropenem | 1.025 | 9.E+05 | 1.E+06 | 12 | 99.125 | 2 log |

#### Imaging infected corneas

All infected and treated corneas were photographed before homogenisation (Figs. 4, 5). *Ex vivo* corneas often swell a little while kept in media for a few days, which may give them a slightly hazy look and affect the opacity grading (uninfected corneas Fig. 4.). Clinically, *P. aeruginosa* keratitis usually manifests with the presence of a large epithelial defect related to stromal necrosis that appears as a ring-like, milky in colour stromal infiltrate. *P. aeruginosa* infection on *ex vivo* porcine cornea manifested with similar features compared to clinical infections *in vivo* (Fig. 4,5 PBS). Visually there was less white discolouration on all corneas treated with MIC concentration of gentamicin, meropenem and ciprofloxacin in comparison to the untreated infected corneas (PBS). This difference is even more obvious on corneas infected with PA14 (Fig. 4B). Corneas infected with both *P. aeruginosa* strains and treated with MIC concentration of gentamicin looked slightly hazier in comparison to meropenem and ciprofloxacin despite no reduction in viable CFU across all antibiotics (Fig. 4).

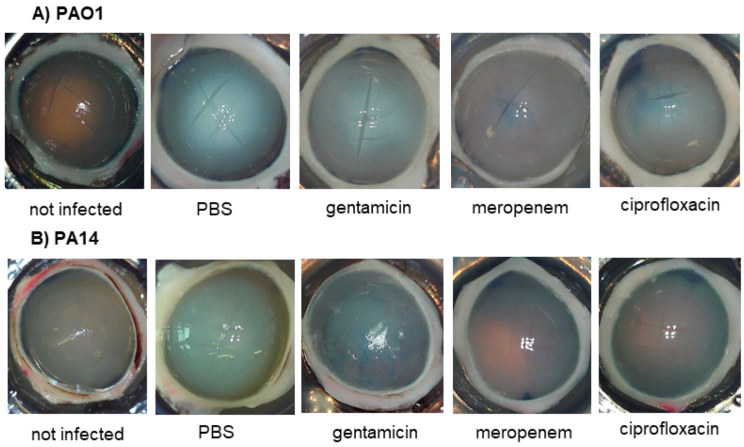


Fig. 4. Representative images of non-infected and infected *ex vivo* porcine corneas without (PBS) treatment and treated with gentamicin, meropenem and ciprofloxacin. The corneas shown here were infected with $6x10^6$ CFU of strain PA01 (A) and strain PA14 (B) and treated with MIC concentrations of antibiotics after the infection had progressed for 6 hours. Corneas were imaged and immediately homogenised for viable counting.

Treatment with a higher antibiotic concentration (1025 μg $mL^{-1}$) decreased corneal opacity of infected cornea by preventing the development of milky colour (Fig. 5). This effect was especially evident in cornea infected with PA14. Corneas treated with gentamicin, meropenem and ciprofloxacin looked clear and visually impossible to distinguish from uninfected showing a direct effect of treatment on opacity (Fig. 5). Despite high bacteria count, gentamicin treatment preserved corneal transparency.

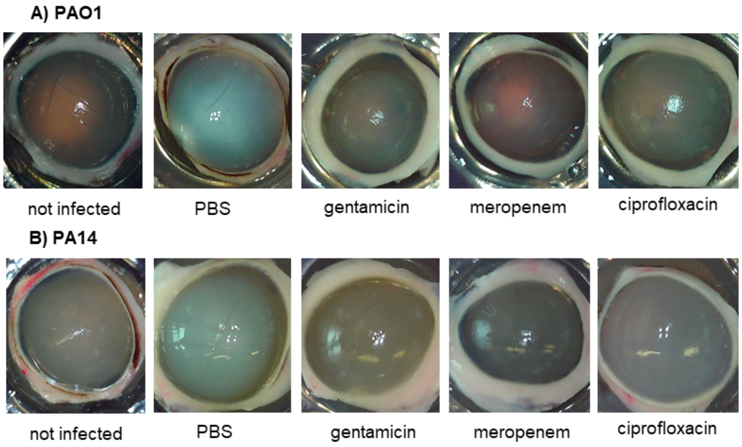


Fig. 5. Representative images of non-infected and infected *ex vivo* porcine corneas without (PBS) treatment and treated with 1025 μg $mL^{-1}$ of gentamicin, ciprofloxacin and chloramphenicol. The corneas shown here were infected with $6x10^6$ CFU of strain PA01 (A) and strain PA14 (B) 6 hours prior to antibiotic treatment. Corneas were imaged and immediately homogenised for viable counting.
